# Attention mechanism-based deep learning pan-specific model for interpretable MHC-I peptide binding prediction

**DOI:** 10.1101/830737

**Authors:** Jing Jin, Zhonghao Liu, Alireza Nasiri, Yuxin Cui, Stephen Louis, Ansi Zhang, Yong Zhao, Jianjun Hu

**Author notes:** Corresponding author, (JH).

## Abstract

Accurate prediction of peptide binding affinity to the major histocompatibility complex (MHC) proteins has the potential to design better therapeutic vaccines. Previous work has shown that pan-specific prediction algorithms can achieve better prediction performance than other approaches. However, most of the top algorithms are neural networks based black box models. Here, we propose DeepAttentionPan, an improved pan-specific model, based on convolutional neural networks and attention mechanisms for more flexible, stable and interpretable MHC-I binding prediction. With the attention mechanism, our ensemble model consisting of 20 trained networks achieves high and more stabilized prediction performance. Extensive tests on IEDB’s weekly benchmark dataset show that our method achieves state-of-the-art prediction performance on 21 test allele datasets. Analysis of the peptide positional attention weights learned by our model demonstrates its capability to capture critical binding positions of the peptides, which leads to mechanistic understanding of MHC-peptide binding with high alignment with experimentally verified results. Furthermore, we show that with transfer learning, our pan model can be fine-tuned for alleles with few samples to achieve additional performance improvement. DeepAttentionPan is freely available as an open source software at https://github.com/jjin49/DeepAttentionPan.

**Author summary:** Human leukocyte antigen (HLA) proteins are classes of proteins that are responsible for immune system regulation in humans. The peptides are short chains of amino acids. HLA class I group present peptides from inside the cell to the cell surface for scrutiny by T cell receptors. For instance, if the cell is infected by a virus, the HLA system will bind to the peptides derived from viral proteins and bring them to the surface of the cell so that the cell can be destroyed by the immune system. Since the HLA genes exhibit extensive polymorphism, there are many HLA alleles binding to different peptides. And this diversity represents challenges in predicting binders for different HLA alleles, which are important in vaccine designs and characterization of immune responses. Before computational algorithms are used to predict the binding relationships of HLA-peptide pairs, scientists need to conduct costly biological experiments to do preliminary screening among a number of peptides and need to use mutant experiments to identify key peptide positions that contribute to the binding. While previous computational methods have been proposed to predict the binding affinity, identifying the binding anchors is not well addressed. Here we developed a deep neural network models with the attention mechanism to learn the binding relationships automatically in an end-to-end way. Our models are able to identify the important binding positions of the peptide sequence by learning the positional importance distribution, which used to be studied a lot only through costly experimental methods. Our model thus not only improves the performance of binding affinity prediction but also allows us to gain biological insight of binding motifs of different alleles via interpreting the learned deep neural network models.

## 1. Introduction

Binding relationships between HLAs and peptides play a critical role in the human immune system. Developing accurate computational methods for their binding affinity prediction has big potential to improve our understanding of the human immune system as well as to help develop novel protein-based drugs [1]. In the past decade, researchers have developed a variety of prediction algorithms for MHC-peptide binding prediction with many successful results [2–7].

There are two major categories of models for MHC-peptide binding affinity prediction: pan-specific models [2, 4, 5, 7–11] such as NetMHCpan [4, 5, 7, 9], DeepSeqPan [10], Pickpocket [8], and ACME [2] and allele-specific models [1, 12–15] such as ANN [13, 14], ARB [16], SMM[12]. Panspecific models are usually trained on datasets of multiple alleles while allele-specific models are developed for specific alleles. Usually allele-specific models can be more accurate when restricted to certain alleles, while pan-specific models deliver more stable and strong overall performances and can be applied to HLA alleles with few samples and even unseen alleles. Due to the wide applications and performance advantages of pan-specific models, the focus of our paper is on developing pan-specific models for MHC peptide binding prediction.

NetMHCpan [9] is the first pan-specific artificial neural network (ANN) model for MHC-peptide binding prediction. The neural network architecture was built based on conventional feed-forward networks with one hidden layer. The network training was performed in a five-fold cross-validation resulting in an ensemble of 25 different independent neural networks. It has shown the advantage of pan-specific methods by exploiting a large number of data across multiple HLA-I supertypes to train a more generalized model with state-of-the-art predicting capability. Its newest version is NetMHCpan 4.0 [4], which is trained on binding affinity and eluted ligand data by leveraging the information from both data types. In the past several years, deep convolutional neural networks (DCNN) with its strong capability in capturing local features and extracting high-level patterns have become very popular in image processing and natural language processing (NLP) [17–20]. Following these successes, DCNN has been applied to MHC-peptide binding prediction in a few recent studies. Kim and Han [11] proposed a DCNN-based pan-specific model and showed the power of DCNN in revealing the locally-clustered patterns in peptide-HLA complexes. Vang and Xie [15] developed a DCNN based allele-specific model with a novel distributed representation of amino acids and has validated the importance of the CNN part in improving the performance. In our previous work, we developed DeepSeqPan [10], which is also a pan-specific model based on two separate DCNN encoders to extract high-level features of both the peptide and the HLA sequence. It is also characterized by a binding context extraction layer using locally connected layers and dual output design. However, most of these algorithms tend to work as black boxes with insufficient interpretation of the learned models to obtain biological insights.

More recently, a pan-specific model, ACME [2], based on DCNN encoders and attention mechanisms has been proposed. It extracts different levels of HLA features from multiple CNN layers and combines them with peptide features separately and sends the concatenated feature vectors to separate fully connected (FC) layers to learn multi-level MHC-peptide relations. It has achieved better performance than many other state-of-the-art methods. Moreover, it is also able to extract interpretable binding motifs based on its attention mechanism [21]. However, there are several major limitations of this model. First, their ablation experiments show that its attention modules have not brought in obvious performance improvements to its base model. The main performance improvement of their algorithm is from the CNN layers. Second, their attention mechanism does not directly encode the position context in addition to their amino acid composition context, which makes it difficult to identify the contribution of different peptide positions to binding. Moreover, the feature map output from their attention module is concatenated to their feature map of the main module just before feeding into the last FC layer of the architecture, which may lead to insufficient pressure needed to train the attention model well. Finally, the final attention map calculated by their formula relies on the 1000 strong binders for each HLA allele that it feeds to the trained models. No results of raw positional weights distribution directly learned by the attention model have been reported for different HLA alleles.

Attention mechanisms [21] are widely used in computer vision (CV) and natural language processing (NLP) [21–23] to help researchers gain insights from trained deep learning models. While previous attempt has achieved some performance improvement by introducing the attention mechanism into MHC-peptide binding prediction[2], here we propose and evaluate three different attention blocks and show that the attention mechanism, when implemented appropriately, can obviously help improve the performance of the model and can provide interpretable positional importance of the peptide in the binding prediction. There are several key features in our attention models. Firstly, our attention module contributes to improving the overall performance of our panspecific models as shown in experiments results (in section 3.4). Secondly, our attention modules learn positional importance of the peptides with different lengths automatically. No extra steps are needed to manually rebuild an attention map as done in ACME [2]. Thirdly, our attention blocks are included in the upper side of the encoders, while the ones in ACME worked separately from the encoders before going into the final FC layers. Our attention modules could work as gates in the encoders directly pushing the model to learn important positions since the weak weights of key positions would directly lead to key information loss. The last but most important feature of our attention module is that it receives an absolute positional context as well as the content information per position. On the contrary, the ACME uses the same single FC layer to learn the attention weights of different positions of a sequence without adding any positional context to the feature map of the sequence. This method to learn attention weights can be used to deal with most content-focused problems, considering that some relative position information could be handled by CNN layers or relative-distance encoding [24]. But in the MHC-peptide binding problems, uncovering the absolute position importance will provide key information about the anchor positions, secondary anchor positions or other effects at certain positions which were experimentally studied by many researchers before [11, 25–28]. Our attention module shows a strong capability to learn related information all together automatically.

In brief, we propose a new DCNN pan-specific model with attention mechanisms for MHC-I peptide binding affinity prediction. It is characterized by its capability to deal with input peptides of variable lengths and its state-of-the-art overall performance. Our attention mechanism improves prediction performance while flexibly capturing important positions of the peptides of different lengths without using any additional encoding techniques to align the peptides. Based on the basic architecture, we use 10-fold cross-validation to train the models and repeat the whole process twice which results in an ensemble of total 20 base models as our final ensemble model, which achieved improved prediction performance and stability. Considering the distinct features of HLA alleles with relatively small number of training samples are more prone to be ignored in pan-specific training due to the simultaneous training over all samples of multiple alleles, we also proposed the idea of transfer learning to fine-tune the pan-model on specific alleles with few training samples, which brings performance improvement for specific alleles by taking advantage of pan-specific models and allele-specific models.

## 2. Materials and methods

### 2.1 Dataset

The training set BD2013 labeled with IC50 binding affinity values is downloaded from IEDB [29]database(http://tools.iedb.org/main/datasets/).

The raw testing set is downloaded from IEDB’s weekly benchmark dataset [1, 29] (http://tools.iedb.org/auto_bench/mhci/weekly/) ranging from 2014-03-21 to 2019-03-15 which is updated to the newest version. Then we delete the duplicate records between training set and raw testing set from the raw testing set to get a preliminary testing set. To be fair, we delete the records in the preliminary testing set that none of the benchmark methods reports predicted values to get the testing set. Furthermore, since the source code of ACME is available online, we have applied a few screening steps mentioned in ACME method to the preliminary testing set to get an additional testing set, the extended testing set. Those screening steps are described in the legend of Table 1. And we compare our algorithm with ACME on the extended testing set rather than the testing set because of more available testing data. All data appearing in the preliminary testing set are not used in model training or model optimization.

**Table 1:** Statistics of the datasets.

| Statistic | Training | Testing | Extended testing |
| --- | --- | --- | --- |
| Number of total samples | 156,844 | 8,855 | 30,554 |
| Number of total binding samples | 40,867 | 4,429 | 19,872 |
| Binding sample percentage | 26.1% | 50.0% | 65.0% |
| Number of HLA alleles | 95 | 25 | 21 |
| Number of datasets | - | 61 | 54 |

The sequences of HLA molecules are downloaded from IMGT/HLA database [30]. More details about the training and testing data are shown in Table 1 below.

The testing set is used to do a benchmark comparison with the IEDB benchmark methods. The extended testing set is only used to make a comparison between our method and ACME [2]. The “datasets” in column label - “Number of datasets”, refer to HLA-peptide sample groups identified by four attributes together - IEDB reference number, HLA allele name, peptide length and measurement type of binding affinity label. And the algorithm performance metrics discussed later are calculated based on every dataset. In particular, the extended testing set has fewer number of alleles and datasets than the testing set since we apply the following screening rules mentioned in ACME to the preliminary testing set to deliver the extended testing set: 1) don’t use datasets with fewer than or equal to 5 samples. 2) only deal with datasets with ic50 or binary labels. Therefore, t1/2 labeled data are simply not used here, considering that t1/2 labeled data is only 7% of the preliminary testing set.

### 2.2 Sequence Encoding

Each input of our model is a raw sequence pair of an HLA allele and a peptide. The HLA-peptide pair is encoded based on the BLOSUM62 scoring matrix. Thus, each residue is encoded by the corresponding row of the scoring matrix with 23 rows and 23 columns. Without loss of generality, in order to receive peptides of different lengths, we set the maximal acceptable peptide length as 15 in our model since most available peptide samples are shorter than 15. Thus, for peptides shorter than 15 residues, we pad zeros at the ends of the sequences to get a 23 ×15 2D tensors. 23 is the number of encoded feature channels. 15 is the length of padded peptide sequences. And in order to receive raw HLA sequences with variable lengths and give our model better tolerance on inputs, we set the maximal acceptable HLA length as 385. For the HLA sequences, we also pad zeros at the ends of the sequences to get 23 × 385 2D tensors.

### 2.3 Architecture of DeepAttentionPan: the attention-based DNN pan-specific model

Figure 1 is the full architecture of our model. It can be divided to two functional submodules. The first submodule is the DCNN main module which contains two DCNN encoders, a binding context extractor, and an affinity predictor. The second submodule is a pair of attention modules. Each is put inside one DCNN encoder following one or a few CNN layers. The outputs of the attention module are fed directly to the rest layers of the DCNN encoder. Below is the detail of each submodule.

**Figure 1:**
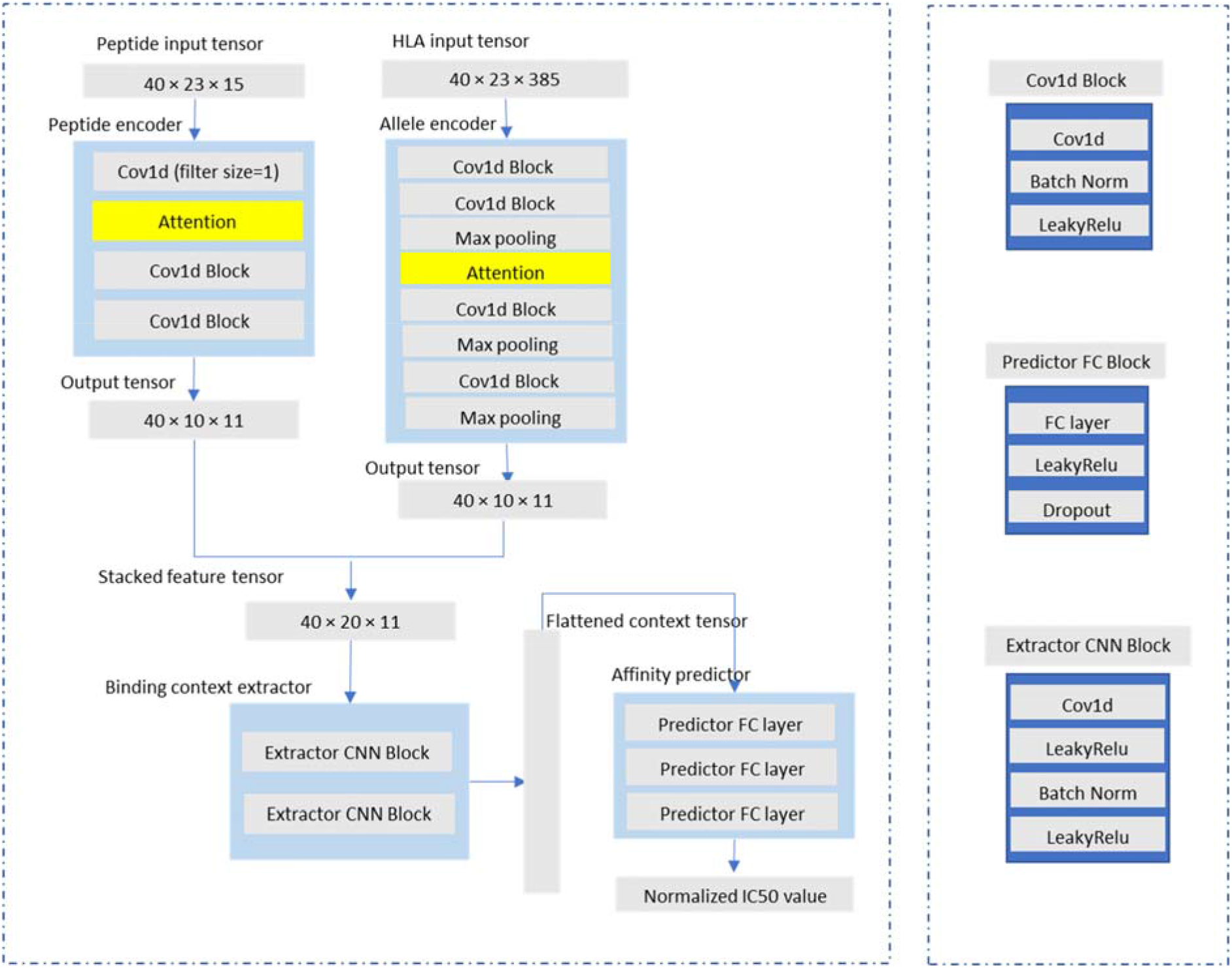
The Architecture of DeepAttentionPan model. The peptide input tensor has a size of 40×23×15 where 40 is the batch size, 23 is the number of encoding channels, and 15 is the maximum sequence length of the peptide. For HLA, the maximum MHC sequence length is 385. The details of the DCNN main module are described in 2.3.1. Attention modules are described in section 2.3.2.

#### 2.3.1 The DCNN main module

The peptide encoder and HLA encoder aim at extracting high-level features of the peptide and the HLA sequence separately. The basic building block of the encoders is a 1D convolutional layer followed by a batch normalization layer and a LeakyReLu activation layer. The encoders convert the peptide and HLA sequences to unified 40 x 10 × 11 vectors where 40 is batch size,10 is the number of the channels of the output sequences, and 11 is the length of output sequences. Then we stack these two vectors via the channel dimension to get a concatenated vector with a dimension of 40 x 20 × 11. Here, the peptide encoder has fewer CNN blocks than the HLA encoder since peptide sequences are shorter. Specifically, we use 1D CNN with a filter size of 1 as the first layer of the Peptide encoder. The filter size used here is important for positional importance learning in the latter attention module since it enables the CNN layer to impose channel-wise transformation to the input sequence as well as to reserve information independence for each position.

Then the binding context extractor will take in the concatenated vector to learn the binding relationship of the peptide-HLA pair and output a binding context vector. It consists of two CNN blocks. In order to add nonlinearity to our model, each CNN block contains one CNN layer, followed by one LeakyRelu layer, a batch-normalized layer and then another LeakyRelu layer. The output binding vector is then flattened to be put into the FC affinity predictor which consists of three FC blocks and outputs the predicted normalized IC50 values at the end. Furthermore, the context extractor helps the model converge better and quicker in the experiments.

#### 2.3.2 Implementation of attention mechanism

In a previous attempt [2] to apply the attention mechanism in MHC-peptide binding, the weight of a position in a sequence is largely based on the content at that position because that a single position-independent FC layer is used to calculate the weights for all positions. In contrast, in our attention mechanism, we add emphasis to the position itself. In other words, the attention module should directly exploit the information of absolute positional difference in weight calculation rather than simply output identical weights for two different positions having identical feature vectors. In fact, our attention mechanisms consider the position and the positional content simultaneously to decide the weights. We will show that, in this way, the model will be able to learn the positional importance of the peptides of different lengths in HLA-peptide binding prediction without any additional manual operations. Considering the efficiency and the purpose, the attention module in each encoder uses different techniques.

For the peptide DCNN encoders, we use multiple independent fully connected (FC) layers (one for each position) to calculate positional weights of the peptide sequence instead of constructing extra explicit position vectors. We name this attention module as ATT1, which is characterized by its implicit position encoding. Figure 2 shows how ATT1 works. We use ATT1 in the peptide encoder because of two reasons. Firstly, peptide raw sequences are much shorter compared to HLA raw sequences, which allows us to use 15 independent FC layers here without affecting efficiency obviously. Secondly, we find this method helps us to capture weights more accurately and flexibly. The model would learn to distinguish different positions automatically as well as considering the contribution of content in each position.

**Figure 2:**
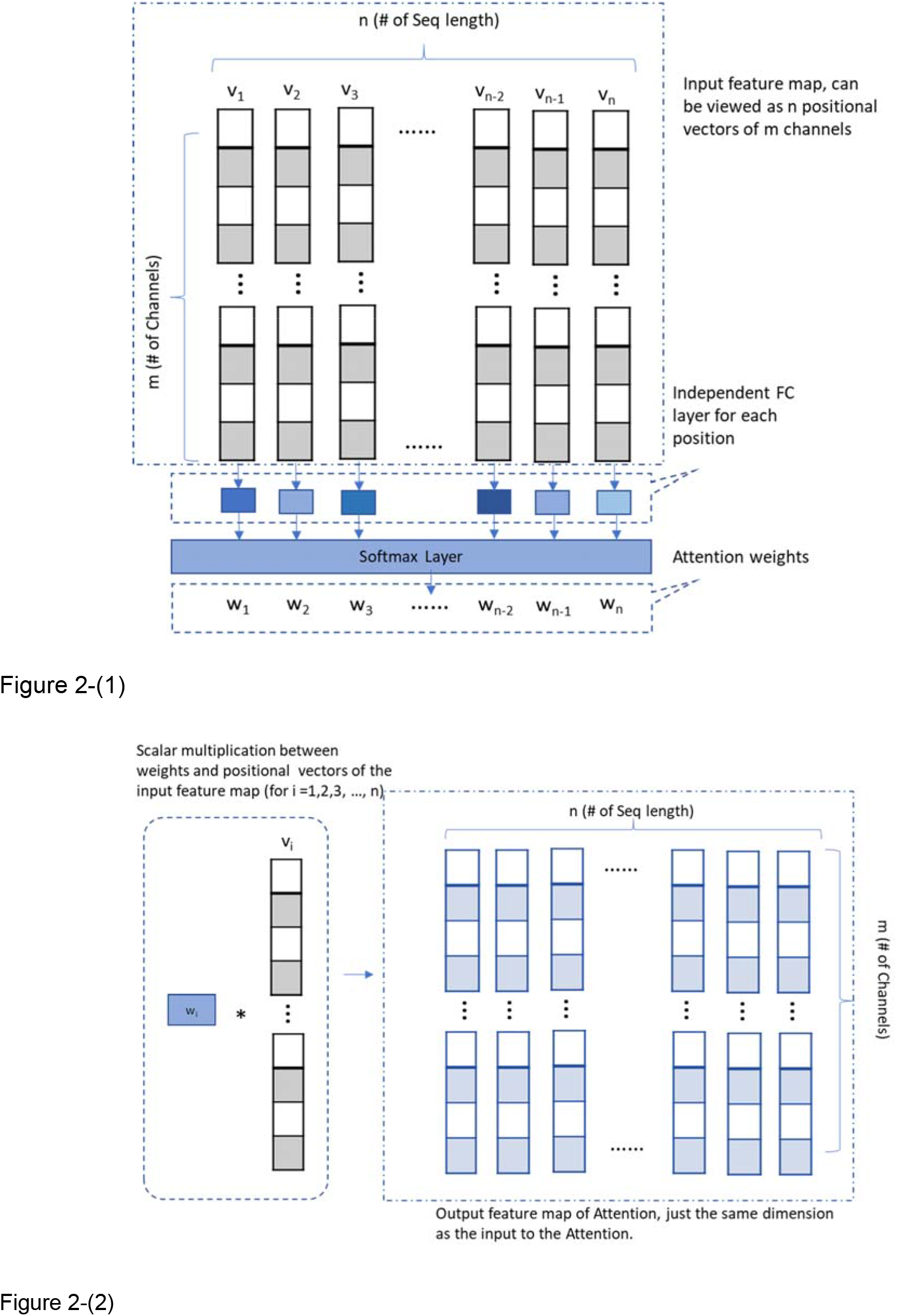
(1) Weight calculation in ATT1 attention module. This part will calculate the attention weights for different positions of the inputs using multiple independent FC layers. (2) Weighted feature map calculation part of ATT1. This part will handle the scalar multiplication between the attention weights and corresponding positional vectors of the input feature map and output a feature map with the same dimension as the input feature map.

For the HLA DCNN encoders, as HLA sequences can be as long as 385 amino acids, it is not feasible or efficient to train 385 independent FC layers for position-dependent weight calculation as done in ATT1. To still achieve position-dependent importance weight calculation, we explicitly encode the positions of the HLA sequence using a one-hot encoding matrix and concatenate it with the HLA feature map coming from the first max-pooling layer of the HLA encoder. And then we use one single FC layer to map the concatenated inputs to calculate the positional weights. This is inspired by the way that relative positional features can be combined with original word feature vectors to build new input used in NLP [31]. We name this attention module as ATT2, which is characterized by its explicit encoding of sequence positions. Figure 3 shows the mechanism of ATT2.

**Figure 3:**
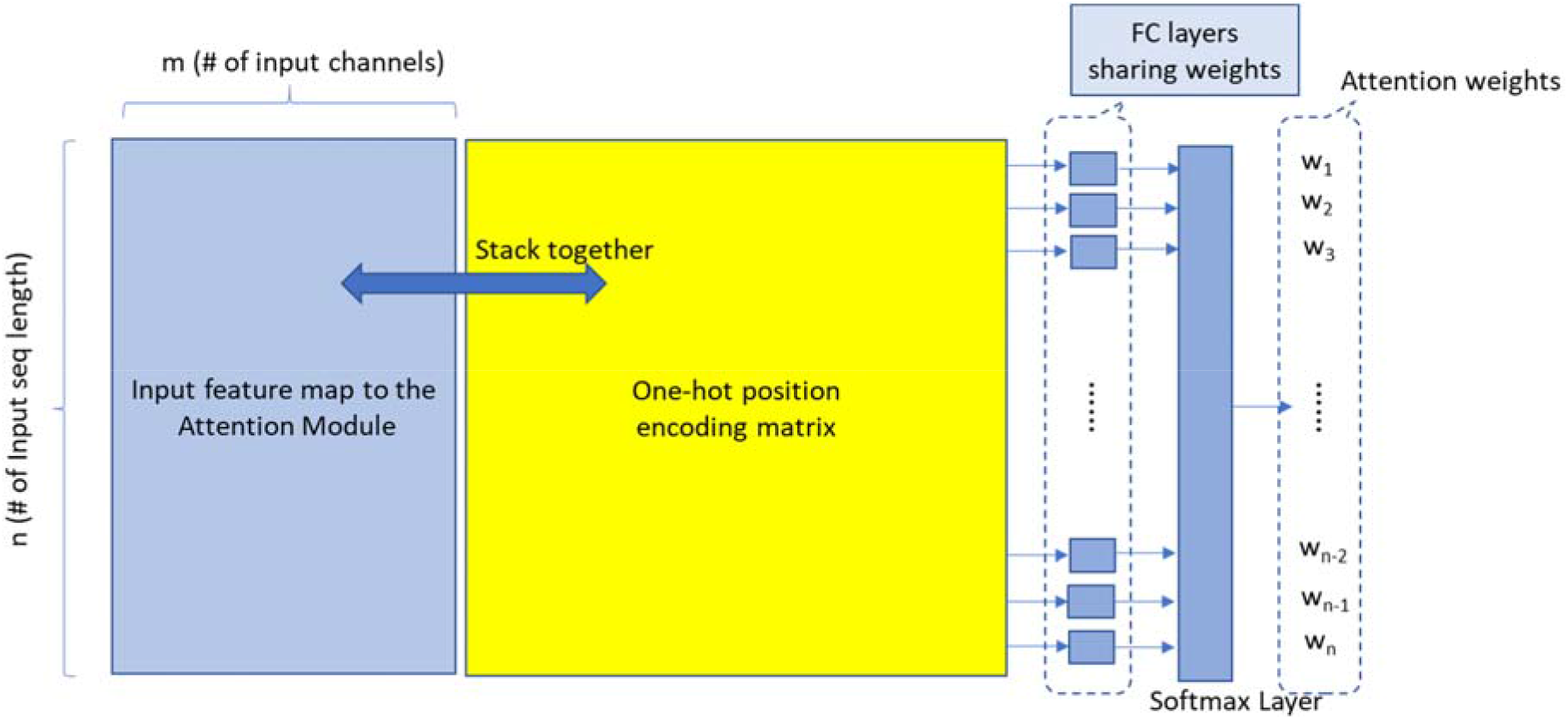
Weight calculation in ATT2 attention module. This part calculates the attention weights based on the inputs of sequences. Weighted feature map calculation of ATT2, which handles the scalar multiplication between the attention weights and the input feature map is the same as the part in ATT1, see Figure 2-(2).

In both ATT1 and ATT2, we use the weights to do scalar multiplication with the original feature vectors and get the output vectors to feed to the next CNN layer in the encoders (see Figure 2-(2)). For ATT2 used in HLA encoder, we first test its effect by applying to the peptide encoder to make sure it can roughly capture the important positions and then apply it to the HLA encoder.

### 2.4 Network training

#### 2.4.1 Training and ensemble

We use a grid search based on 10-cross validation to tune the hyperparameters of our prediction models. When we train our final deep neural network models, we use a batch size of 40, SGD optimizer with a reducible learning rate starting at 0.2. Our final model is an ensemble of 20 networks from two consecutive 10-cross validation experiments.

#### 2.4.2 Metrics and label preprocessing

AUC (Area under the curve) is used as the main evaluation metric, and SRCC (Spearman’s rank correlation coefficient) is used as an additional evaluation metric. Because of the sparse distribution of IC50 values, the training label of the HLA-peptide pair is normalized as score = 1- (log(IC_50_)/log(50000)). We use the normalized label in our training and then convert the prediction results back to the original IC_50_ values in the testing. In our experiments, a HLA-peptide pair with its original IC_50_ value less than 500 nM is a binding pair.

### 2.5 Transfer learning

We propose to exploit the idea of transfer learning [32] to build better allele-specific models by taking advantage of pan-specific models. While pan-specific algorithms can exploit larger training set to build more stable models, the number of training samples for different HLA alleles are very unbalanced. Therefore, the unique features of some HLA alleles with less training samples may be ignored in the training process as the model could be dominated by several HLA alleles with much more training samples. On the one hand, we want to preserve the stability of the panspecific models. On the other hand, we want to improve the performance of the prediction models for the minority. This leads us to fine-tune our pan-specific models with the training samples of distinct alleles separately.

The total number of training samples (including the validation sets) we use is 156,844. The number of corresponding alleles is 95. Therefore, the average number of samples per allele is 1650. We select the alleles with less than 1650 and more than 100 samples. And we fine-tune our model on those specific alleles separately, using their own training samples. For each allele, we use a very small starting learning rate of 0.005 to train 20 new networks with initial parameters loaded from the 20 base networks of the trained pan model. Then we output the ensemble model of the 20 new networks and test it on the testing samples of the target allele. As we show in section 4, just with this small learning rate, we are able to bring stable improvements over the pan model through transfer learning. Furthermore, for all the experiment results listed in section 3, transfer learning is not involved to improve the models.

## 3. Results

### 3.1 Performance Evaluation on benchmark datasets

To compare our pan-specific model with other HLA-peptide binding algorithms, we use IEDB weekly benchmark dataset [1] as our raw testing set (ranging from 2014-03-21 to 2019-03-15 and is updated to the newest version), on which some widely-known algorithms have published their prediction results. The duplicate records of the training set and raw testing set are removed from the raw testing set. Samples in the raw testing set that all other benchmark methods do not report prediction values are also deleted in order to achieve more fair comparison. Then we get our final testing set, in short, the testing set to conduct the benchmark comparison. We have compared the performance of our model with these models: ‘NetMHCpan 2.8’ [7], ‘NetMHCpan 3.0’[5], ‘NetMHCpan 4.0’ [4], ‘SMM’ [12], ‘ANN 3.4’ [13], ‘ANN 4.0’ [14], ‘ARB’ [1], ‘SMMPMBEC’ [1], ‘IEDB Consensus’ [1], ‘NetMHCcons’ [6], ‘PickPocket’ [8].

Table 2 summarizes the performance of all the algorithms above on the 61 datasets of the testing set. The last row of Table 2 is the number of the datasets upon which each algorithm achieves the highest scores. And our model and NetMHCpan4.0 both achieve the largest number of datasets upon which the target model has the highest AUC scores, which is 21. In terms of SRCC, NetMHCpan4.0 ranks first on 19 datasets, which is the best. Our model and ANN4.0 rank second both with 17 datasets. However, after a close examination, we obtain a more clear understanding when our algorithm will outperform others.

**Table 2:**
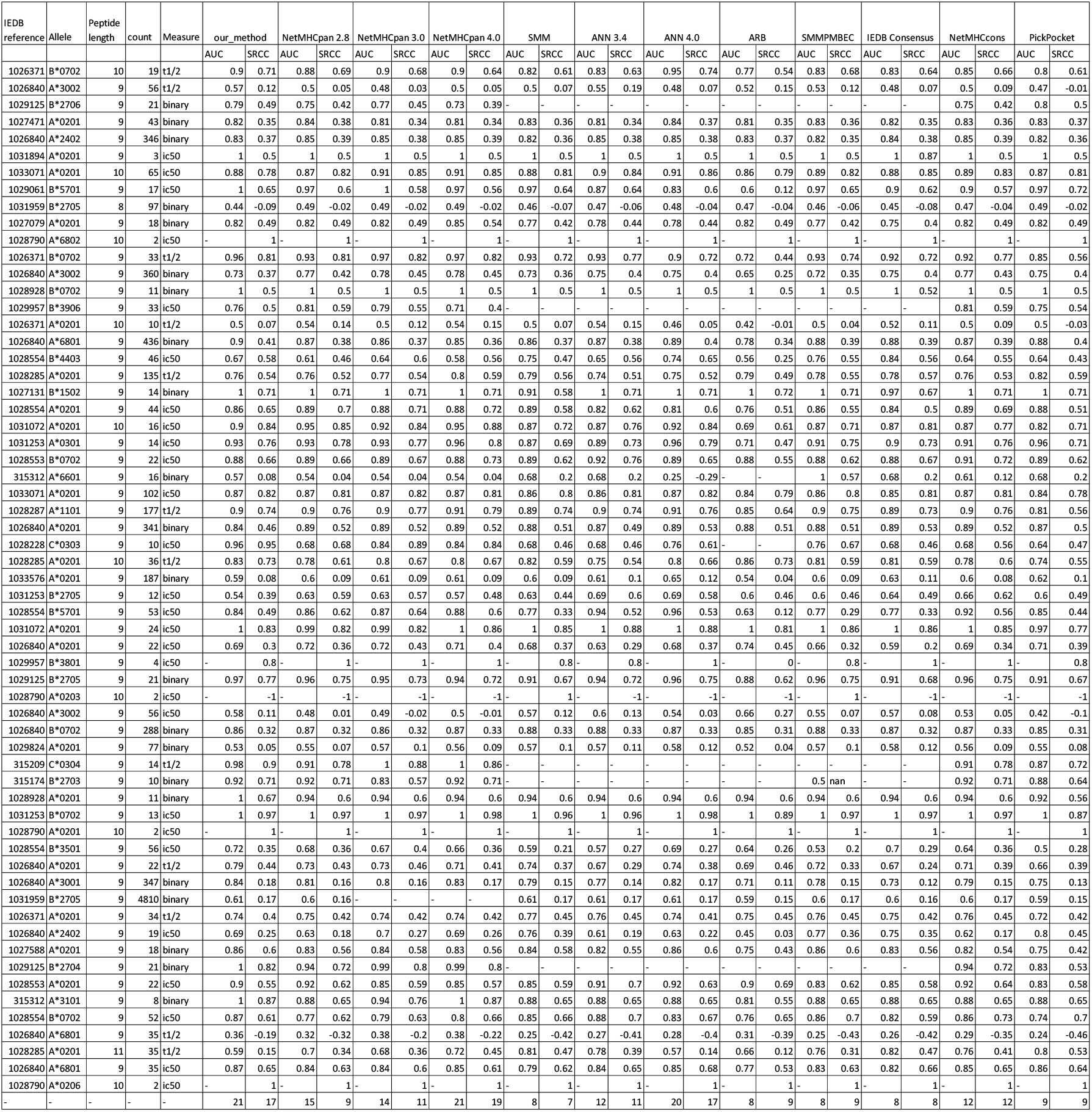
performance of our model against benchmark algorithms on all 61 datasets. The last row of the Table is the number of the datasets upon which each algorithm achieves the highest scores.

To gain more insight into the prediction characteristics of our algorithm, we compare our model only with NetMHCpan4.0 on the testing set. The allele-specific comparing results are shown in Figure 4 based on the average AUC scores and in Figure 5 based on average SRCC scores. There are 4 HLA alleles in the testing set in Table 2 that both methods don’t have AUC scores due to small numbers of samples. Therefore, both methods are compared on the remaining 21 HLA alleles.

**Figure 4:**
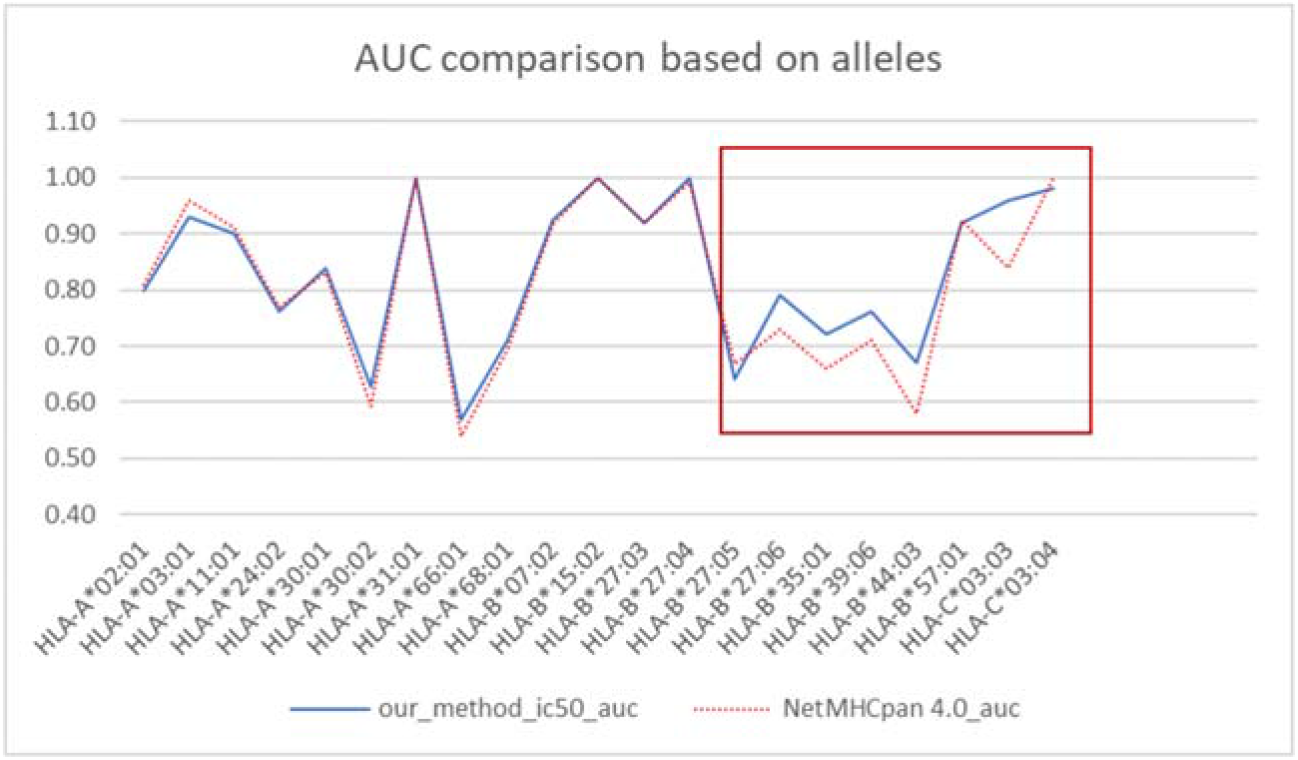
AUC comparison between our model and NetMHCpan4.0. The highlighted box area shows that our model achieves obvious better performance for several alleles than NetMHCpan4.0. There are 10 alleles on which our model has AUC scores at least 0.01 higher than NetMHCpan4.0’s and only on 5 alleles that scores of NetMHCpan4.0 are at least 0.01 higher than ours.

**Figure 5:**
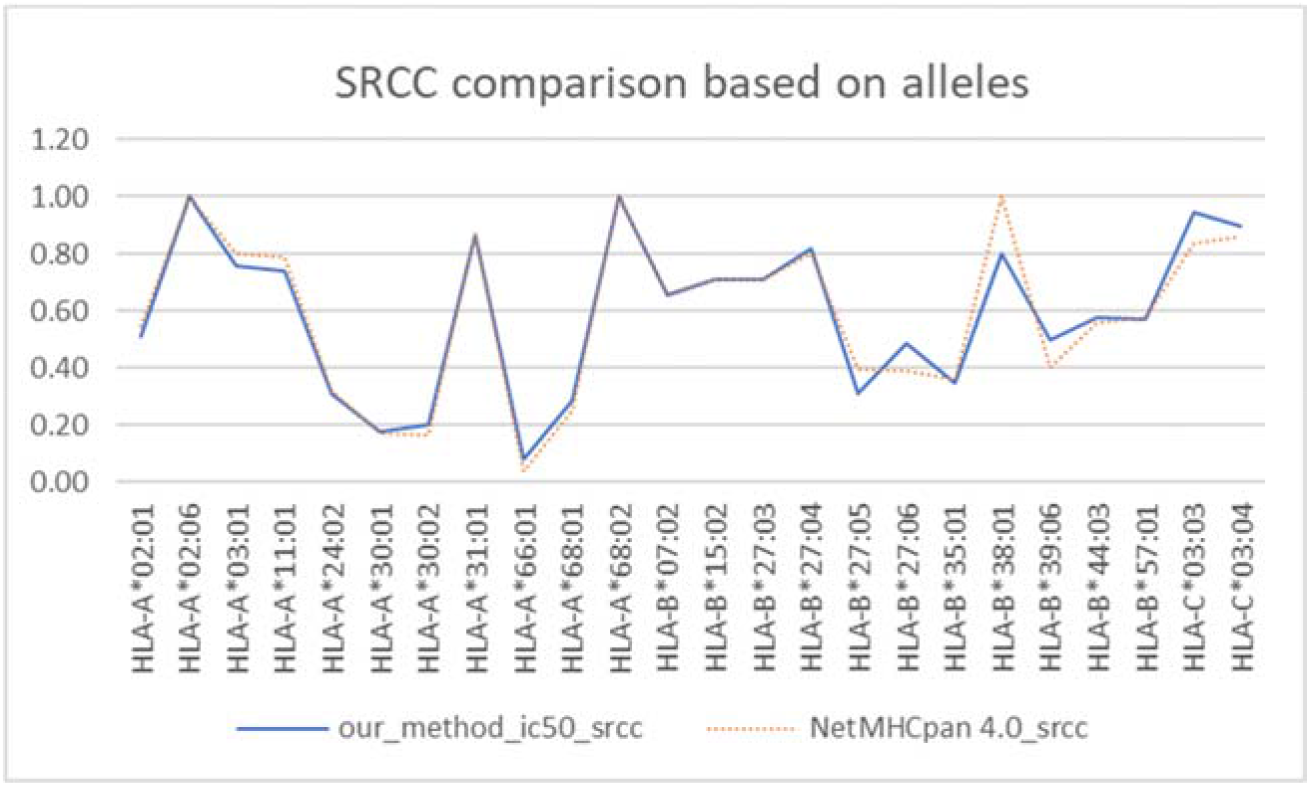
SRCC comparison between our model and NetMHCpan4.0. There are 9 alleles on which our scores are at least 0.01 higher than NetMHCpan4.0’s and vice versa.

In Figure 4, there are several alleles on which our model achieves obvious better performance in average AUC scores. We put a box in Figure 4 to highlight those alleles. And for other HLA alleles, our model also delivers comparable results to NetMHCpan4.0. In summary, there are 10 alleles for which our model has scores at least 0.01 higher than NetMHCpan4.0. On the contrary, there are only 5 alleles for which the scores of NetMHCpan4.0 are at least 0.01 higher than ours. So, at the allele level, our DeepAttentionPan algorithm has overall better performance. Moreover, Table 3 below shows that our model usually has an advantage on the HLA alleles with few training samples. And this makes sense since our attention module can help stabilize the training process and achieve better generalization performance by focusing on important positions and their information.

**Table 3:** The number of samples of the HLA-alleles grouped by the “winning method with AUC metric”. Our method has an obvious advantage over the alleles with fewer training samples.

| **NetMHCpan4.0 is at least 0.01 higher in AUC** | **Training samples** | **Testing samples** |
| --- | --- | --- |
| HLA-A*03:01 | 7358 | 14 |
| HLA-A*11:01 | 6399 | 177 |
| HLA-A*24:02 | 3280 | 365 |
| HLA-B*27:05 | 3676 | 4940 |
| HLA-C*03:04 | 0 | 14 |
| **DeepAttentionPan is at least 0.01 higher in AUC** | **Training samples** | **Testing samples** |
| HLA-A*30:01 | 2925 | 347 |
| HLA-A*30:02 | 2057 | 472 |
| HLA-A*66:01 | 219 | 16 |
| HLA-A*68:01 | 3795 | 506 |
| HLA-B*27:04 | 6 | 21 |
| HLA-B*27:06 | 7 | 21 |
| HLA-B*35:01 | 3211 | 56 |
| HLA-B*39:06 | 0 | 33 |
| HLA-B*44:03 | 1404 | 46 |
| HLA-C*03:03 | 154 | 10 |

In Figure 5, in terms of average SRCC score, NetMHCpan4.0 outperforms our algorithm obviously on HLA-B*38:01 and HLA-B*27:05. But our model shows obvious advantage on HLA-B*27:06, HLA-B*39:06 and HLA-C*03:03. Moreover, despite that we don’t have any training samples related to HLA-B*39:06 or HLA-C*03:03, our model still has good prediction results to those two alleles in the testing set. In sum, there are 9 alleles for which our SRCC scores are at least 0.01 higher than NetMHCpan4.0 and 9 alleles for which the scores of NetMHCpan4.0 are at least 0.01 higher than ours. Therefore, two models are just as good in terms of SRCC scores.

We have stored the more detailed comparison results on all 61 datasets in supplementary file 1. In short, when we only consider the two methods, our model and NetMHCpan4.0 both have 34 datasets that rank the first in terms of AUC scores. In terms of SRCC, our model achieves the highest scores on 32 datasets compared to 38 of NetMHCpan4.0. However, at the allele level, our model shows overall better performance considering both AUC scores and SRCC scores. As our algorithm demonstrates a clear advantage over alleles with few or unknown binding peptides, our method is a good complementary model to the NetMHCpan4.0. Moreover, our algorithm only uses the IC50 binding values with HLA-peptide sequence pairs to train the model, while the NetMHCpan4.0 also takes auxiliary eluted ligand datasets. Overall, from all the results above, our model shows an impressive prediction capability compared to other benchmark algorithms.

### 3.2 Performance comparison with ACME on extended testing set

We make an extensive and fair comparison of our method with the ACME, another deep learning-based prediction model also equipped with the attention modules [2], over the extended testing set which is also built from the IEDB weekly benchmark dataset [1]but with more data than the testing set used above. This is possible as ACME is open-sourced, unlike NetMHCPan4.0. ACME is a newly developed method that has achieved state-of-the-art prediction results. The detail about the extended testing set is shown in section 2.1. We use the published code of ACME and set its training set identical to ours and then test both methods on the same extended testing set. We run the experiments for ACME for five times. Each time it delivers an ensemble model of 25 networks. Then we choose the ensemble model with the best overall testing result to compare with our model, which is the same as they did in their evaluation experiments.

Table 4 below shows the comparison results. There are 23 datasets upon which our model has higher AUC scores than ACME and only 15 datasets upon which ACME has higher AUC scores. In terms of SRCC, there are 24 datasets upon which we have higher scores than ACME and 22 datasets upon which ACME has higher scores than ours.

**Table 4:** Comparison results between DeepAttentionPan and ACME. Red ones refer to the higher scores. There are 23 datasets upon which we have higher AUC scores than ACME and only 15 datasets upon which ACME has higher AUC scores than us. In terms of SRCC, there are 24 datasets upon which we have higher scores than ACME and 22 datasets upon which ACME has higher scores than us. In the allele-specific level, our model also achieves better results than ACME both in AUC and SRCC. Details are in section 3.2.

| Dataset | Allele | Peptide Length | Measure | Count | ACME SRCC | OUR SRCC | ACME AUC | OUR AUC |
| --- | --- | --- | --- | --- | --- | --- | --- | --- |
| 1033576 | A*02:01 | 9 | binary | 187 | 0.04 | 0.08 | 0.54 | 0.59 |
| 1031072 | A*02:01 | 9 | ic50 | 24 | 0.86 | 0.83 | 1 | 1 |
| 1031072 | A*02:01 | 10 | ic50 | 16 | 0.83 | 0.84 | 0.87 | 0.9 |
| 1033071 | A*02:01 | 9 | ic50 | 102 | 0.81 | 0.82 | 0.86 | 0.87 |
| 1033071 | A*02:01 | 10 | ic50 | 65 | 0.79 | 0.78 | 0.91 | 0.88 |
| 1031959 | B*27:05 | 8 | binary | 97 | -0.1 | -0.09 | 0.44 | 0.44 |
| 1031959 | B*27:05 | 9 | binary | 13540 | 0.16 | 0.17 | 0.6 | 0.6 |
| 1031959 | B*27:05 | 10 | binary | 7663 | 0.2 | 0.19 | 0.63 | 0.63 |
| 1031959 | B*27:05 | 11 | binary | 5330 | 0.24 | 0.23 | 0.66 | 0.66 |
| 1031253 | A*03:01 | 9 | ic50 | 14 | 0.79 | 0.76 | 0.91 | 0.93 |
| 1031253 | B*07:02 | 9 | ic50 | 13 | 0.95 | 0.97 | 1 | 1 |
| 1031253 | B*27:05 | 9 | ic50 | 12 | 0.63 | 0.39 | 0.66 | 0.54 |
| 315312 | A*66:01 | 9 | binary | 16 | 0.29 | 0.08 | 0.75 | 0.57 |
| 315312 | A*31:01 | 9 | binary | 8 | 0.87 | 0.87 | 1 | 1 |
| 1029957 | B*39:06 | 9 | ic50 | 33 | 0.57 | 0.5 | 0.77 | 0.76 |
| 1029824 | A*02:01 | 9 | binary | 77 | 0.12 | 0.05 | 0.58 | 0.53 |
| 1027131 | B*15:02 | 9 | binary | 14 | 0.71 | 0.71 | 1 | 1 |
| 1029125 | B*27:06 | 9 | binary | 21 | 0.4 | 0.49 | 0.74 | 0.79 |
| 1029125 | B*27:04 | 9 | binary | 21 | 0.78 | 0.82 | 0.98 | 1 |
| 1029125 | B*27:05 | 9 | binary | 21 | 0.72 | 0.77 | 0.94 | 0.97 |
| 1029061 | B*57:01 | 9 | ic50 | 17 | 0.66 | 0.65 | 1 | 1 |
| 1028928 | A*02:01 | 9 | binary | 11 | 0.6 | 0.67 | 0.94 | 1 |
| 1028928 | B*07:02 | 9 | binary | 11 | 0.5 | 0.5 | 1 | 1 |
| 315174 | B*27:03 | 9 | binary | 10 | 0.57 | 0.71 | 0.83 | 0.92 |
| 1028553 | A*02:01 | 9 | ic50 | 22 | 0.58 | 0.55 | 0.92 | 0.9 |
| 1028553 | B*07:02 | 9 | ic50 | 22 | 0.63 | 0.66 | 0.93 | 0.88 |
| 1028554 | B*57:01 | 9 | ic50 | 53 | 0.54 | 0.49 | 0.88 | 0.84 |
| 1028554 | B*44:03 | 9 | ic50 | 46 | 0.62 | 0.58 | 0.67 | 0.67 |
| 1028554 | B*35:01 | 9 | ic50 | 56 | 0.33 | 0.35 | 0.58 | 0.72 |
| 1028554 | A*02:01 | 9 | ic50 | 44 | 0.76 | 0.65 | 0.92 | 0.86 |
| 1028554 | B*07:02 | 9 | ic50 | 52 | 0.62 | 0.61 | 0.84 | 0.87 |
| 1027588 | A*02:01 | 9 | binary | 18 | 0.52 | 0.6 | 0.81 | 0.86 |
| 1027471 | A*02:01 | 9 | binary | 43 | 0.36 | 0.35 | 0.82 | 0.82 |
| 1027079 | A*02:01 | 9 | binary | 18 | 0.54 | 0.49 | 0.85 | 0.82 |
| 1026897 | B*40:01 | 9 | binary | 15 | 0.38 | 0.38 | 0.75 | 0.75 |
| 1026897 | B*40:01 | 10 | binary | 11 | 0.5 | 0.5 | 1 | 1 |
| 1026897 | B*58:01 | 9 | binary | 22 | 0.3 | 0.32 | 0.75 | 0.77 |
| 1026897 | B*58:01 | 10 | binary | 18 | 0.3 | 0.39 | 0.73 | 0.8 |
| 1026891 | A*11:01 | 9 | binary | 16 | 0 | -0.12 | 0.5 | 0.39 |
| 1026891 | A*24:02 | 9 | binary | 19 | -0.05 | 0.26 | 0.47 | 0.68 |
| 1026891 | B*40:01 | 9 | binary | 19 | 0.58 | 0.77 | 0.83 | 0.94 |
| 1026891 | B*58:01 | 9 | binary | 20 | 0.62 | 0.62 | 0.86 | 0.86 |
| 1026840 | A*24:02 | 9 | binary | 346 | 0.37 | 0.37 | 0.83 | 0.83 |
| 1026840 | A*24:02 | 9 | ic50 | 19 | 0.08 | 0.25 | 0.58 | 0.69 |
| 1026840 | A*30:01 | 9 | binary | 347 | 0.15 | 0.18 | 0.8 | 0.84 |
| 1026840 | B*58:01 | 9 | binary | 433 | 0.4 | 0.43 | 0.88 | 0.91 |
| 1026840 | B*58:01 | 9 | ic50 | 34 | 0.2 | 0.32 | 0.62 | 0.7 |
| 1026840 | B*07:02 | 9 | binary | 288 | 0.33 | 0.32 | 0.87 | 0.86 |
| 1026840 | A*30:02 | 9 | binary | 360 | 0.4 | 0.37 | 0.75 | 0.73 |
| 1026840 | A*30:02 | 9 | ic50 | 56 | 0.05 | 0.11 | 0.55 | 0.58 |
| 1026840 | A*68:01 | 9 | binary | 436 | 0.41 | 0.41 | 0.89 | 0.9 |
| 1026840 | A*68:01 | 9 | ic50 | 35 | 0.58 | 0.65 | 0.79 | 0.87 |
| 1026840 | A*02:01 | 9 | binary | 341 | 0.47 | 0.46 | 0.85 | 0.84 |
| 1026840 | A*02:01 | 9 | ic50 | 22 | 0.4 | 0.3 | 0.72 | 0.69 |

We also conduct allele-specific comparison. There are 10 alleles over which our AUC scores are at least 0.01 higher than ACME’s and only over 5 alleles that ACME achieves AUC scores at least 0.01 higher than us. In terms of SRCC, it is 10 (our model) compared to 8 (ACME). Therefore, from both dataset level and allele level, our model achieves better performance on the extended testing set than ACME.

### 3.3 Biological insight extracted by the attention mechanism

Attention mechanisms in our deep neural network provide a way to interpret the trained neural network models by calculating the importance or contributions of the amino acid positions of the peptides for MHC-peptide binding. To verify if our attention module can rediscover the important positions identified in previous experimental studies, we use the following procedure to check. After training our final model, we fix the parameters, feed different groups of samples into the model to get the average positional weights of those peptide groups.

Firstly, we use the samples in the testing set and group them by the peptide length and feed them to the trained pan-specific model. We have calculated the average positional weight distribution for three different length groups (length_9, length_10, length_11). The result is shown in Figure 6. It shows that either position 2 or the C-terminus get the highest attention weight for each group and both positions’ weights always rank the top three. The model’s capability of dynamically capturing the importance of the C-terminus is shown clearly on the figure by the curves’ rightmost peaks, which automatically move one position to the right as the length increase by one unit. This is in accordance with the previous experimental studies [25–27] since position 2 and the C-terminus are two primary anchor positions for most alleles. Moreover, position 1 also has high weights. This matches the results of previous experimental studies [26, 28], which point out position 1 is among the secondary anchor positions for many MHC alleles. The results above demonstrate the capability and potential of our attention modules to uncover biological mechanisms in MHC-peptide binding.

**Figure 6:**
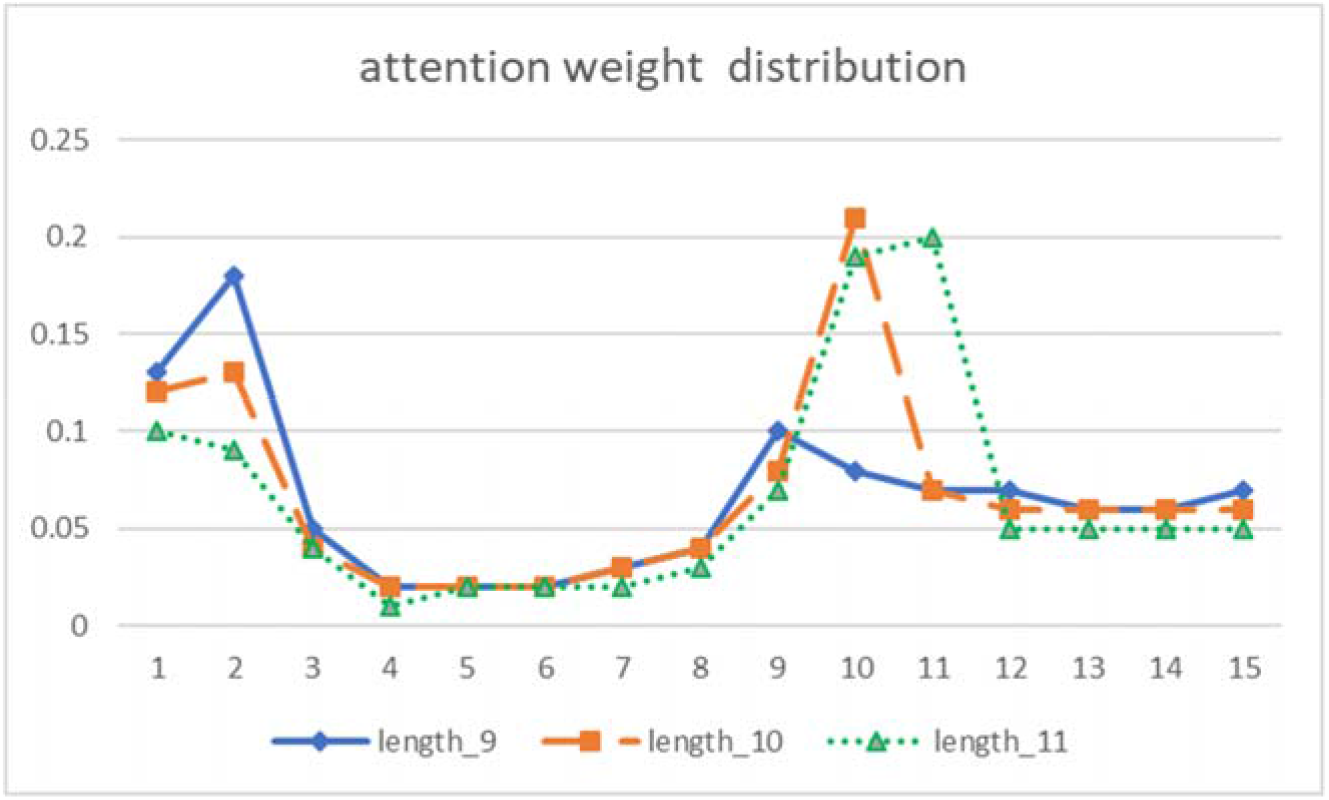
Attention weight distributions grouped by peptide length. The importance of position 2 in binding is captured by the left peaks of the curves. And the importance of the C-terminus is shown by the dynamic movements of curves’ right-most peaks. Position 1 is also assigned with a relatively high weight by our algorithm, which is the secondary anchor position for many HLA alleles [26, 28].

And then, we use the 9-length binding samples from the training set and group them by each HLA allele. This is for allele-specific experiments. Since the attention module in our pan-specific model is trained to deliver the positional importance distribution of peptides for the majority of H LA alleles, we found it is not flexible enough to show extraordinary anchors of a minority of HLA alleles. In order to get more exact peptide weight distributions for specific alleles to investigate peptide-binding positional importance, we choose to train a group of allele-specific models using the same architecture and training process, while each training set is restricted to samples of the target allele. The picked HLA alleles all have more than 1000 training samples and most of the alleles are analyzed in the experimental study [26]. The normalized weight distributions without considering the padding positions of different alleles are listed in Table 5 below. The results from [26] as one control group are shown in the column labeled “PSCLA”. “PSCLA” is short for the Positional Scanning Combinatorial Library Approach [26]. And the positional conservation distributions of peptides are used as another control group. The conservation level per position for a HLA allele is calculated by score = 5 – (–Σ_*i*_(*p_i_*log*pi*)). The *p_i_* is the probability of the *i^th^* amino acid out of 20 amino acids appearing at this position in all 9-length binders related to certain HLA allele. From the Table 5, the attention maps show obvious weight disparity among different positions with several highlights deserve our attention.

**Table 5:** Positional weight distributions of peptides for different HLA alleles. The attention weight distributions are compared with the experimental PSCLA method [26] and positional amino acid conservation distributions, which is also used to validate our attention module. The attention module shows its advantage than other two control groups in terms of anchor identification by integrating different information to decide the importance of each position to binding. It is able to assign lower attention weights to non-anchor positions despite their high amino acid conservation (see HLA*B-2705). It can distinguish secondary anchor positions with non-important positions despite their similar amino acid conservation levels.

| Alleles<br>(Sample<br>size) | Normalized attention weight | Conservation distribution | PSCLA[12] | Alleles<br>(Sample<br>size) | Normalized attention weight | Conservation distribution | PSCLA[12] |
| --- | --- | --- | --- | --- | --- | --- | --- |
| A*0201<br>(9051) |  |  | P1,P2,P3,P9 | B*2705<br>(2811) |  |  | P2 |
| A*3001<br>(2565) |  |  | P1,P2,P3,P9 | B*3501<br>(2514) |  |  | P2,P9 |
| A*6802<br>(3672) |  |  | P1,P2,P3,P9 | B*5101<br>(2239) |  |  | P2,P9 |
| B*0702<br>(3868) |  |  | P2,P3,P9 | B*5301<br>(1052) |  |  | P1,P2,P3,P9 |
| B*0801<br>(3027) |  |  | P1,P2,P3,P5,P6,P9 | B*5801<br>(2984) |  |  | P1,P2,P7,P9 |
| B*1501<br>(4101) |  |  | P2,P9 | A*0301<br>(5488) |  |  | - |

First of all, our attention module has a strong capability to pick out the main anchor positions. The primary anchor positions detected in the conservation distribution are always assigned with high weights by our models.

Typical instances include HLA-B*0801 and HLA-A*3001. Both alleles have distinct primary anchor positions from the majority. In the case of HLA-B*0801, the positions 3, 5, and the C-terminus are identified as primary anchor positions in [33]. And all three primary anchor positions of HLA-B*0801 have high attention weights, especially the positions 3 and 5, which are unusual anchors. However, position 3 is not detected as a primary anchor position by “PSCLA”, which may due to lack of binding data. In the case of HLA-A*3001, position 3 is one of the primary anchors in the experimental study [26], which also shows a high value in its conservation level. And in our model, position 3 is assigned the third highest weight, obviously higher than the non-anchor positions.

Another example is HLA-B*2705, for which the main anchor is position 2 without the C-terminus as confirmed in the experimental study [20, 28]. Our attention module here assigns much higher attention weight to position 2 than others, which is consistent with the experimental result. Furthermore, even the C-terminus shows obviously higher conservation, our model doesn’t assign equivalent importance to this position, which shows its capability to integrate even more diverse information than positional conservation in its learning process.

Secondly, our attention module shows strong potential to distinguish the secondary anchors with non-important positions and recognize other locally-clustering properties, which may not be reflected in amino acid conservation distributions. A good example is allele HLA-A*0201 with abundant training samples. The secondary anchor positions are experimentally identified as 1,3,7 in [28]. We can distinguish these three positions much more clearly in the attention map rather than in the conservation distribution. This is significant since even the “PSCLA” experimental technique, as well as conservation distribution, cannot recognize position 7 as a secondary anchor. In another case study of the HLA-A*0301 allele, our model assigns high weights to both position 1 and position 2 which is consistent with the experimental study mentioned in [11] showing that there exist locally-clustering interactions at the two positions of the binding complex, which help stabilize the binding, thus confirming our prediction.

### 3.4 Comparison of performance of three different attention mechanisms

In order to study the prediction performance improvement and capability of capturing important positions bringing by position-dependent attention mechanisms, we implement three different attention modules in the peptide encoder of the pan-specific model. We analyze their resulting weight distributions and compare their final prediction performance. We group the testing set by peptide length to fed to the trained models to deliver the average positional weights of peptides the same as the first experiment conducted in section 3.3.

ATT1 and ATT2 refer to the corresponding attention modules as we defined in section 2.3.2. We have named the attention module with multiple independent FC layers as ATT1 and the attention module with a single FC layer but with concatenated inputs with encoded position matrices as ATT2. Attention weight distribution on peptides is shown in Figure 6 for Att1 and in Figure 7-(a) for ATT2. We define ATT3 as the control group which refers to the attention module with a single position-independent FC layer without using any positional matrix, which is similar to what ACME algorithm does. ATT3’s weight distribution on peptides is shown in Figure 7-(b). From Figure 7-(a), ATT2 is able to learn that two ends of the peptides are important positions. But it is not flexible enough to capture the real C-terminus of peptides with different lengths: the rightmost peaks all locate at position 10 due to apparent averaging effect. In Figure 7-(b), ATT3 is only able to distinguish padding positions with the original sequence in terms of attention weights. It totally fails to distinguish any two different positions in the original sequence in terms of their importance or contribution to binding. This clearly demonstrates the advantages of our position-dependent attention mechanism.

**Figure 7:**
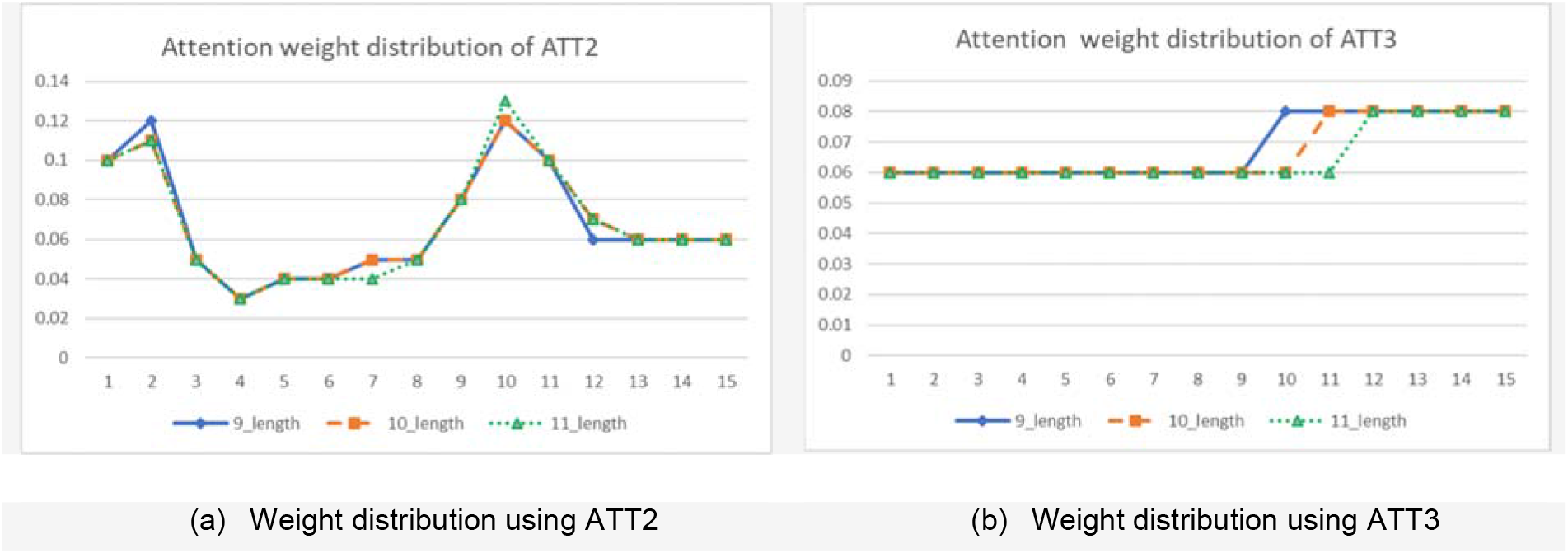
Attention weight distribution grouped by different peptide length Comparison of different attention mechanisms for modeling the importance of peptide positions in MHC-I binding. ATT2 is the attention module using a single FC layer with combined inputs of feature map and positional matrix. It can roughly capture the importance of two ends of the peptide. But it can’t dynamically capture the C-terminus of peptides of different lengths as ATT1 in Figure 6 does. ATT3 is the control group that refers to the attention module using a single position-independent FC layer without using any positional matrix. It totally fails to distinguish the important difference of the positions in original sequences.

Furthermore, we have compared the influences that different attention modules bring to the prediction capability of our models. We are comparing the results based on the overall prediction performance scores on the testing set in Table 6. For each architecture, we run 5 replicated experiments. Each experiment delivers an ensemble model of 20 trained networks using 10 crossvalidation method. We also list the average scores and standard deviations of five experiments for each architecture in Table 6.

**Table 6:** This is the overall performance comparison among three attention methods on the testing set. ATT1 and ATT2 both have improved the prediction performance from the baseline in both average scores and standard deviations. And ATT1 has achieved the best performance by all metrics listed in the table. The ATT3 doesn’t work better than the baseline in terms of the average scores.

| Experiment | ATT1<br>AUC | ATT1<br>SRCC | ATT2<br>AUC | ATT2<br>SRCC | ATT3<br>AUC | ATT3<br>SRCC | NO_ATT<br>AUC | NO_ATT<br>SRCC |
| --- | --- | --- | --- | --- | --- | --- | --- | --- |
| 1 | 0.807 | 0.505 | 0.801 | 0.490 | 0.791 | 0.487 | 0.794 | 0.490 |
| 2 | 0.808 | 0.507 | 0.803 | 0.501 | 0.788 | 0.491 | 0.788 | 0.482 |
| 3 | 0.809 | 0.511 | 0.801 | 0.494 | 0.791 | 0.486 | 0.792 | 0.491 |
| 4 | 0.805 | 0.506 | 0.801 | 0.495 | 0.784 | 0.482 | 0.792 | 0.483 |
| 5 | 0.808 | 0.503 | 0.801 | 0.492 | 0.788 | 0.482 | 0.791 | 0.499 |
| Average | **0.808** | **0.506** | 0.801 | 0.495 | 0.788 | 0.485 | 0.791 | 0.489 |
| Std | **0.001** | **0.002** | 0.001 | 0.004 | 0.002 | 0.003 | 0.002 | 0.006 |

The “NO_ATT” in Table 6 refers to the architecture removing both attention modules in encoders, which is used as the baseline. Considering the average scores of 5 experiments, ATT3 does not bring any performance improvement from the baseline. ATT2 and ATT1 improve from the baseline in both average scores and standard deviation scores. And Att1 Ranks first by all the metrics listed below, which is 0.808 in average AUC, 0.506 in average SRCC, 0.001 in standard deviation of AUC, and 0.002 in standard deviation of SRCC. It has obviously improved the overall performance as well as the stability of the training results. The overall performance increase from baseline to ATT1 is 0.016 in AUC and 0.017 in SRCC.

## 4. Discussion

### 4.1 Effect of transfer learning

While the pan-specific model can capture the overall binding patterns, we also check if the general model can be fine-tuned to achieve better prediction performance on a given allele using the idea of transfer learning. Here we show the results of transfer learning on all 6 alleles which have fewer training samples than average in Table 7. In order to make the comparison more convincible, we adopt two control groups. One is our final pan-specific model, another one is the allele-specific model built in the totally same way as the pan model except for the training set. For each target HLA allele, we repeat the experiments of transfer learning and allele-specific training for 5 times each in order to deliver statistically significant results. And we calculate the average scores and standard deviations of repeated experiments to check the average performance and stability.

**Table 7:** Performance improvements with transfer learning. We compared the results from transfer learning with our pan-specific model and the corresponding allele-specific models. The scores of transfer learning and the allele-specific models both are the average scores of 5 repeated experiments. The results show that transfer learning achieves the highest average scores on every HLA allele in the table and its standard deviations of five experiments are much lower than the allele-specific models. The performance improvements transferring from the pan models are relatively stable according to these results.

| Alleles | Training set size | Pan model |  | Transferred model |  |  |  | Allele-specific model |  |  |  |
| --- | --- | --- | --- | --- | --- | --- | --- | --- | --- | --- | --- |
|  |  | Average |  | Average |  | Std |  | Average |  | Std |  |
|  |  | AUC | SRCC | AUC | SRCC | AUC | SRCC | AUC | SRCC | AUC | SRCC |
| B*44:03 | 1404 | 0.67 | 0.58 | **0.70** | **0.62** | 0.004 | 0.007 | 0.68 | 0.60 | 0.0040 | 0.0172 |
| B*27:03 | 876 | 0.92 | 0.71 | 0.92 | 0.71 | 0.025 | 0.044 | 0.51 | 0.01 | 0.1851 | 0.3166 |
| B*38:01 | 517 | - | 0.8 | - | **1.00** | - | 0.000 | - | 0.60 | - | 0.1789 |
| A*66:01 | 219 | 0.57 | 0.08 | **0.69** | **0.21** | 0.012 | 0.020 | 0.67 | 0.19 | 0.1666 | 0.1911 |
| B*15:02 | 164 | 1 | 0.71 | 1.00 | 0.71 | 0.000 | 0.000 | 0.77 | 0.38 | 0.1145 | 0.1643 |
| C*03:03 | 154 | 0.96 | 0.95 | **1.00** | **0.96** | 0.000 | 0.004 | 0.66 | 0.53 | 0.0480 | 0.0882 |
| Average |  | 0.82 | 0.64 | **0.86** | **0.70** | 0.008 | 0.013 | 0.66 | 0.38 | 0.1036 | 0.1594 |

From the results in Table 7, the average scores of transfer learning for each allele are always ranking the first among the three methods. And it shows its performance stability from much smaller standard deviations, which is 0.008 and 0.013 on average, than those of allele-specific models, which is 0.1036 and 0.1549 on average. The results of allele-specific models are unstable due to a very small number of training samples. While adopting pan-specific model largely decreases the performance instability because of a large training set. Based on this fact, model transferring from the pan model with a small learning rate is able to preserve the good stability of pan-specific models, as well as able to focus on the unique features of specific alleles to improve performance. In all, the average performance of the transferred models is 0.86 in AUC and 0.70 in SRCC, compared to 0.82 and 0.64 of the pan model, 0.66 and 0.38 of allele-specific models. Therefore, transfer learning shows promising improvements from two control groups. In short, transfer learning may work as a bridge between the pan-specific model and allele-specific model to improve performance on the alleles with few training samples.

### 4.2 The importance of ensemble

In our experiment, we use the ensemble model consisting of 20 networks trained from the same architecture to deliver the final model. This is because the ensemble method helps improve the binding affinity prediction performances steadily and results in more robust models. The average AUC score of our final ensemble model is 0.81 over 61 datasets, compared to 0.78 of the best single network from 20 independent experiments. In terms of the average SRCC score, the ensemble model is 0.5, and the single best model is 0.48. The average scores of both metrics have obviously improved. In terms of testing stability of the ensemble model, we conduct 5 experiments and find the overall scores remain very stable among 5 experiments with an average AUC of 0.81 and an average SRCC of 0.51. The corresponding standard deviation scores are 0.001 and 0.002, which is very small. The details of the five repeated experiments are in Table 6.

## 5. Conclusion

We propose a pan-specific MHC binding prediction model based on deep neural networks with attention mechanisms. This model is characterized by its performance stability by taking advantage of the ensemble of networks, flexibility to deal with input peptide sequences with different lengths and the interpretability brought by the attention modules, which can be conveniently implemented and added to other MHC-peptide binding prediction algorithm architectures. We have also conducted experiments using transfer learning to combine the advantage of pan-specific models and the allele-specific models.

Extensive evaluation of our pan-specific model on the public benchmark experiments shows that our model achieves state-of-the-art performance on diverse HLA allele datasets. The comparison experiment of DeepAttentionPan with ACME shows the overall advantage of our model on the extended testing set. Detailed analysis and interpretation of our attention module shows that it is able to capture important binding-contributing positions of the peptides of varying lengths, which has been verified by previous experimental studies. Additional experiments corresponding to each HLA allele further showed the capabilities of the attention module to integrate diverse information useful for improving binding affinity prediction. It is expected that our three different ways of attention mechanisms in binding prediction can also be widely applied to deep learning models for other biological problems such as protein-protein interaction, protein-RNA/DNA binding and etc.

## Author Contributions

Conceived and designed the experiments: JH, JJ; Performed the experiments: JJ ZL; Analyzed the data: JJ JH ZL; Wrote the paper: JJ JH. Provided methodological input: ZL YC YZ AN.

## Notes

#### Summary of Updates

Title updated. Figures updated to latest version.

https://github.com/jjin49/DeepAttentionPan

